# Phosphatidylinositol 5 phosphate 4 kinase regulates phosphatidylinositol 3,4 bisphosphate levels *in vivo*

**DOI:** 10.1101/2025.11.24.690171

**Authors:** Aishwarya Venugopal, Shreya Varma, Devaprabha Varsha, Raghu Padinjat

## Abstract

In *Drosophila*, loss of phosphatidylinositol 5 phosphate 4 kinase (PIP4K) results in a reduction in larval salivary gland cell size. Previous studies have shown that this reduction in cell size is not correlated with the levels of phosphatidylinositol 5 phosphate (PI5P), the canonical substrate of PIP4K but to the levels of phosphatidylinositol 3 phosphate (PI3P) a substrate that is used less effectively by the PIP4K enzyme *in vitro*. The phosphorylation of PI3P by PIP4K generates phosphatidylinositol 3,4 bisphosphate [PI(3,4)P_2_]. Using a biosensor for PI(3,4)P_2_, surprisingly, we find that depletion of PIP4K leads to an elevation of intracellular PI(3,4)P_2_ punctae in salivary gland cells. This elevation in PI(3,4)P_2_ punctae was not dependent on the catalytic activity of dPIP4K. Rather, we found that the elevation of PI(3,4)P_2_ was dependent on the catalytic activity of Class II phosphatidylinositol 3 kinase (Class II PI3K). Thus, the PIP4K protein regulates an intracellular pool of PI(3,4)P_2_ via Class II PI3K activity in *Drosophila* cells.

## Introduction

The response of cells to changes in the environment is mediated by communication mechanisms that involve the use of chemical messengers which convey information across the plasma membrane. Cell surface receptors that detect extracellular stimuli lead to the generation of chemical signals within cells, that tune cellular biochemistry to meet ongoing needs. One such class of chemical signals generated in cells are phosphoinositides, generated by differential phosphorylation of the hydroxyl group at the 3^rd^, 4^th^, and 5^th^ positions on the inositol head group. This combinatorial modification produces a family of seven phosphoinositides, namely three mono-phosphorylated regioisomers PI3P, phosphatidylinositol 4 phosphate (PI4P) and PI5P, three bis-phosphorylated regioisomers - PI(4,5)P_2_, PI(3,4)P_2_ and PI(3,5)P_2_ and one tris-phosphorylated isomer- PI(3,4,5)P_3_ (Posor et al., 2022). Each of the species functions as a discrete second messenger in eukaryotic cells. Individual phosphoinositides bind specifically to cellular proteins thereby relaying information during cell signalling.

Phosphatidylinositol (3,4)-bisphosphate [PI(3,4)P_2_] is one of the phosphoinositides whose resting levels in cells is very low [(Ray et al., 2024);(Dickson & Hille, 2019)]. The majority of cellular PI(3,4)P_2_ is produced downstream of receptor mediated PI3K activation in which phosphatidylinositol (3,4,5)-trisphosphate [PIP₃] serves as the precursor for PI(3,4)P₂ [(Hawkins et al., 1992);(Stephens et al., 1991)]. PI(3,4)P_2_ was for long considered as a minor and inconsequential by-product of PIP_3_ metabolism. However, a growing body of evidence has demonstrated that PI(3,4)P_2_ itself performs signalling functions. It regulates critical cellular and physiological processes such as clathrin mediated endocytosis (CME)(Posor et al., 2013), metabolism (Dong et al., 2019) and tumour metastasis [(Gewinner et al., 2009);(S. Ghosh et al., 2018)].

As with all signalling molecules the ability of PI(3,4)P_2_ to control cellular physiology depends on its generation in a precise spatial and temporal profile. This itself is controlled by the activity of precisely localized lipid kinases and phosphatases that receive upstream signals and are activated to add or remove phosphate groups (Balla, 2013) from the inositol head group. Thus, the activity of the lipid kinases and phosphatases that control PI(3,4)P_2_ is central to its signalling functions in cells. Two pathways primarily responsible for PI(3,4)P_2_ production have been reported. The first involves the sequential action of a 3-kinase (Class I PI3K) on phosphatidylinositol (4,5)-bisphosphate [PI(4,5)P_2_] to convert it into PIP_3_ and the subsequent action of a 5-phosphatase (e.g., SHIP1/2) to convert PIP_3_ to PI(3,4)P_2_ [(Damen et al., 1996);(Pesesse et al., 1997)].This mechanism predominantly generates PI(3,4)P_2_ at the plasma membrane. A recent study has suggested that this pool of PI(3,4)P_2_ can be endocytosed via CME to give rise to PI(3,4)P_2_ at early endosomes (Liu et al., 2018). An alternate pathway for PI(3,4)P_2_ production is regulated by the Class II PI3Ks. The mammalian genome contains three genes encoding these large multidomain enzymes (PI3K-C2α, PI3K-C2β, and PI3K-C2γ) [(Domin et al., 1997),(Rozycka et al., 1998) (Misawa et al., 1998)] whereas less complex metazoans such as *Drosophila* have only one gene i.e., Pi3K68D (MacDougall et al., 2004). Class II PI3Ks can synthesize both PI3P by phosphorylation of PI and PI(3,4)P_2_ by phosphorylation of PI4P *in vivo* [(Braccini et al., 2015);(Velichkova et al., 2010)]. Recently, much attention has focused on the cellular functions of PI(3,4)P_2_ produced from PI4P by class II PI3K [(Marat et al., 2017);(Posor et al., 2013)]. On the other hand, PI(3,4)P_2_ can be metabolised by selective degradation through inositol polyphosphate 4-phosphatases (INPP4A and INPP4B) [(Norris & Majerus, 1994);(Norris et al., 1995);(Gewinner et al., 2009)] or the 3-phosphatase (PTEN) [(Malek et al., 2017);(Goulden et al., 2019)].

The first indication of a PI3P to PI(3,4)P_2_ conversion pathway came from studies in Swiss 3T3 fibroblasts, where oxidative stress induced selective PI(3,4)P_2_ accumulation alongside a proportional decrease in PI3P, pointing to the involvement of a PI3P-specific 4-kinase (Van Der Kaay et al., 1999). The critical gap that still remained at the time was the identification of the kinase responsible for this activity. Subsequent work implicated phosphatidylinositol 4 phosphate 5 kinase (PIP4K) in this process. PIP4K is a metazoan specific lipid kinase(Krishnan et al., 2025) that efficiently phosphorylates PI5P and generates PI(4,5)P_2_. Consistent with this, experiments in multiple species and cells has shown that depletion of PIP4K leads to an increase in PI5P levels [reviewed in (Krishnan et al., 2026)]. Zhang et al. (Zhang et al., 1997) showed that PIP4K could convert PI3P into PI(3,4)P_2_, a finding further confirmed by Rameh et al (Rameh et al., 1997). Complementary studies in mammalian cells showed that double knockdown of PIP4K2A/B reduced PI(3,4)P_2_ levels (Emerling et al., 2013), whereas overexpression of PIP4K2B increased PI(3,4)P_2_ in the context of p110Caax expression(Carricaburu et al., 2003), thus reinforcing the idea of a direct conversion of PI3P to PI(3,4)P_2_. A study in *Drosophila* has also reported that PI3P levels are elevated in flies depleted of PIP4K in a kinase dependent manner (A. Ghosh et al., 2023). However, the ability of PIP4K to directly phosphorylate PI3P to generate PI(3,4)P_2_ in cells remains to be unequivocally established.

In this study, we have developed tools to determine the levels and distribution of PI(3,4)P_2_ in *Drosophila* cells. For this purpose, we have used the C-terminal PH domain of TAPP1 (Dowler et al., 2000) protein repeated thrice in tandem fused to eGFP (eGFP::cPHx3), a high avidity probe for PI(3,4)P_2_ detection, as developed by Goulden et al (Goulden et al., 2019). Using this probe, we find that *Drosophila* cells show both plasma membrane and endomembrane pools of PI(3,4)P_2_. We find that the endomembrane pool of PI(3,4)P_2_ is virtually undetectable in wild type cells but accumulates at high levels in cells depleted of dPIP4K. Interestingly this elevation of PI(3,4)P_2_ levels could be rescued by a kinase dead version of dPIP4K indicating a non-catalytic mechanism of regulation of PI(3,4)P_2_ levels. Further, we find that the elevated levels of PI(3,4)P_2_ likely arises from the catalytic activity of a Class II PI3K driven pathway. Thus, our studies define a role for dPIP4K in the control of a Class II PI3K dependent pool of PI(3,4)P_2_ in *Drosophila* cells.

## Results

### eGFP::cPHx3 reports levels of PI(3,4)P_2_ in *Drosophila* cells

We expressed the full-length TAPP1::eGFP and the eGFP::cPHx3 probe in *Drosophila* S2R+ cells **[Supplementary Fig 1A]**. The eGFP::cPHx3 probe is predominantly enriched at the plasma membrane with very few punctate structures as compared to the full length TAPP1::eGFP probe **[Supplementary Fig 1B (a and b) and 1C]**. By contrast, a non-binding version of the eGFP::cPHx3 probe (eGFP::R211LcPHx3) **[Supplementary Fig 1A]** in which a critical residue Arginine (R) is mutated to Leucine (L) in all the three cPH domains, is diffused throughout the cytosol **[Supplementary Fig 1B (c) and 1C].** This finding highlights that the probe specifically reports PI(3,4)P_2_ distribution. Notably, the TAPP1-based probe detected PI(3,4)P₂ at levels comparable to those reported by the non-binding eGFP::R211LcPHx3 probe, indicating that the low avidity of TAPP1 probe severely limits its use for detecting endogenous levels of PI(3,4)P₂.

To explore the responsiveness of the eGFP::cPHx3 probe to changes in PI(3,4)P_2_ levels, we starved S2R+ cells. In principle, starvation should reduce PIP_3_ levels (Schwarzer et al., 2006) and thus as a consequence reduce PI(3,4)P_2_ on the plasma membrane. As expected, starving cells for 30 mins decreased plasma membrane associated eGFP::cPHx3 fluorescence but did not alter the distribution of the non-binding probe eGFP::R211LcPHx3 [**Fig 1, A and B**].

**Figure 1:**
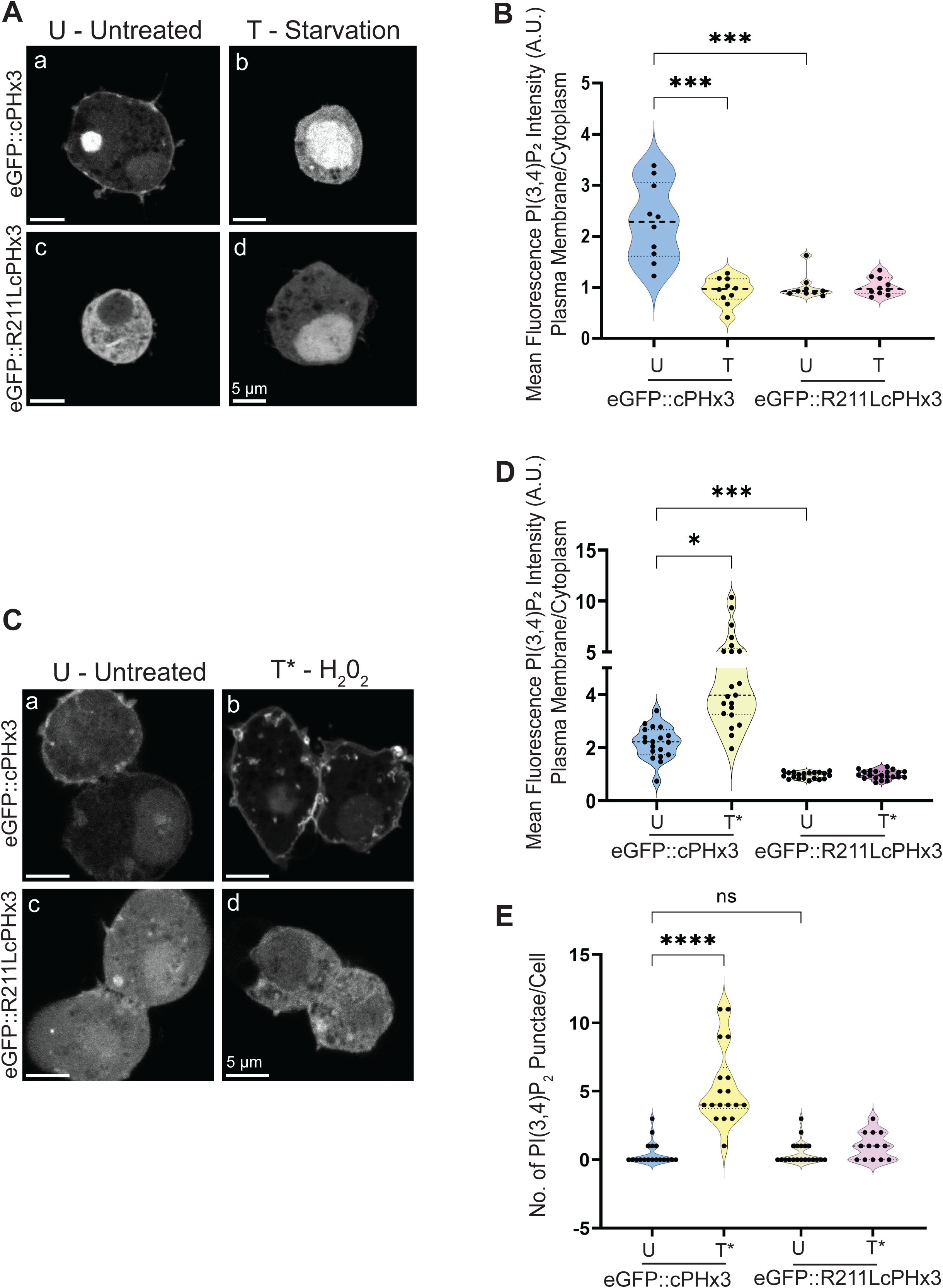
eGFP::cPHx3 reports levels of PI(3,4)P_2_ in *Drosophila* cells. **(A)** Representative confocal z-projections illustrating PI(3,4)P_2_ levels using eGFP::cPHx3 and eGFP::R211LcPHx3 in the S2R+ cells from the following genotypes: (a) U - *Act>eGFP::cPHx3* Untreated, (b) T - *Act>eGFP::cPHx3* Starvation treated, (c) U - *Act>eGFP::R211LcPHx3* Untreated and (d) T *- Act> eGFP::R211LcPHx3* Starvation treated. The scale bar is indicated at 5 μm for spatial reference. **(B)** Quantification of PI(3,4)P_2_ mean fluorescence intensity at the plasma membrane using eGFP::cPHx3 and eGFP::R211LcPHx3 in the S2R+ cells from the following genotypes: (a) U - *Act>eGFP::cPHx3* Untreated (n=10), (b) T- *Act>eGFP::cPHx3* Starvation treated (n=10), (c) U - *Act>eGFP::R211LcPHx3* Untreated (n=9) and (d) T - *Act>eGFP::R211LcPHx3*. Starvation treated (n=10). Statistical test: Kruskal-Wallis Test (***P value < 0.001). **(C)** Representative confocal z-projections illustrating PI(3,4)P_2_ levels using eGFP::cPHx3 and eGFP::R211LcPHx3 in the S2R+ cells from the following genotypes: (a) U - *Act>eGFP::cPHx3* Untreated, (b) T* - *Act>eGFP::cPHx3* H_2_O_2_ treated, (c) U - *Act> eGFP::R211LcPHx3* Untreated and (d) T* - *Act> eGFP::R211LcPHx3* H_2_O_2_ treated. The scale bar is indicated at 5 μm for spatial reference. **(D)** Quantification of PI(3,4)P_2_ mean fluorescence intensity at the plasma membrane using eGFP::cPHx3 and eGFP::R211LcPHx3 in the S2R+ cells from the following genotypes: (a) U - *Act>eGFP::cPHx3* Untreated (n=19), (b) T* - *Act>eGFP::cPHx3* H_2_O_2_ treated (n=21), (c) U - *Act>eGFP::R211LcPHx3* Untreated (n=20) and (d) T* - *Act>eGFP::R211LcPHx3* H_2_O_2_ treated (n=21). Statistical test: Kruskal-Wallis Test (*P value = 0.019, ***P value < 0.001). **(E)** Quantification of endomembranous PI(3,4)P_2_ levels using eGFP::cPHx3 probe and eGFP::R211LcPHx3 probe in the S2R+ cells from the following genotypes:(a) *Act>eGFP::cPHx3* Untreated (n=18), (b) *Act>eGFP::cPHx3* H_2_O_2_ treated (n=18), (c) *Act> eGFP::R211LcPHx3* Untreated (n=20) and (d) *Act>eGFP::R211LcPHx3* H_2_O_2_ treated (n=13). Statistical test: Kruskal-Wallis Test (****P value<0.0001, ns – P value>0.9999).

Hydrogen peroxide (H_2_O_2_) stimulation is known to increase plasma membrane PI(3,4)P_2_ levels (Dowler et al., 2000). On treating cells with H_2_O_2_, we observed a robust increase of eGFP::cPHx3 on the plasma membrane [**Fig 1, C and D**] along with an increase in discrete eGFP::cPHx3 decorated endomembranous compartments [**Fig 1, C and E**]. These punctae were positive for the early endosomal protein Rabenosyn-5 (data not shown) suggesting that the punctae most likely originated from the plasma membrane through endocytosis. The increase in eGFP::cPHx3 intensity at the plasma membrane and punctae were specific to PI(3,4)P_2_ as the distribution and intensity of eGFP::R211LcPHx3 did not increase in response to H_2_O_2_ treatment [**Fig 1, D and E**]. Together, these findings establish the robustness of the eGFP::cPHx3 probe in reporting both plasma membrane as well as the endomembranous pools of PI(3,4)P_2_ in *Drosophila* cells. To carry out further *in vivo* experiments, we generated transgenic flies expressing eGFP::cPHx3 and eGFP::R211LcPHx3.

### dPIP4K regulates PI(3,4)P_2_ levels in *Drosophila* salivary gland cells

We expressed eGFP::cPHx3 in *Drosophila* larval salivary gland cells and observed no obvious localization of the probe to either the plasma membrane or endomembranous structures **[Fig 2A (a)]**. To test the ability of dPIP4K to convert PI3P to PI(3,4)P_2_, we overexpressed dPIP4K in salivary glands but found no increase in PI(3,4)P_2_ levels **[Supplementary Fig 2, A and B]**. On the other hand, in *dPIP4K^29^* (loss of function allele), we observed that PI(3,4)P_2_ levels were strongly upregulated in the salivary glands **[Fig 2, A and B]**. Immunoblotting of salivary gland extracts showed equivalent expression of eGFP::cPHx3 between control and *dPIP4K^29^* confirming that variations in probe expression was not the underlying cause for elevation in the number of PI(3,4)P_2_ punctae in *dPIP4K^29^* **[Supplementary Fig 2C]**. Our finding of increased PI(3,4)P_2_ punctae in *dPIP4K^29^* was recapitulated on RNA interference (RNAi) mediated dPIP4K downregulation in salivary glands **[Fig 2, C and D]** despite equivalent levels of probe expression in control and RNAi glands **[Supplementary Fig 2D]**. The punctae were specific to PI(3,4)P_2_ as a non-binding version of the probe (eGFP::R211LcPHx3) failed to pick up any PI(3,4)P_2_ punctate signal **[Supplementary Fig 3, B and C];** expression levels of both the binding and non-binding probes were comparable across genotypes, ruling out probe expression differences as a confounding factor **[Supplementary Fig 3A]**.

**Figure 2:**
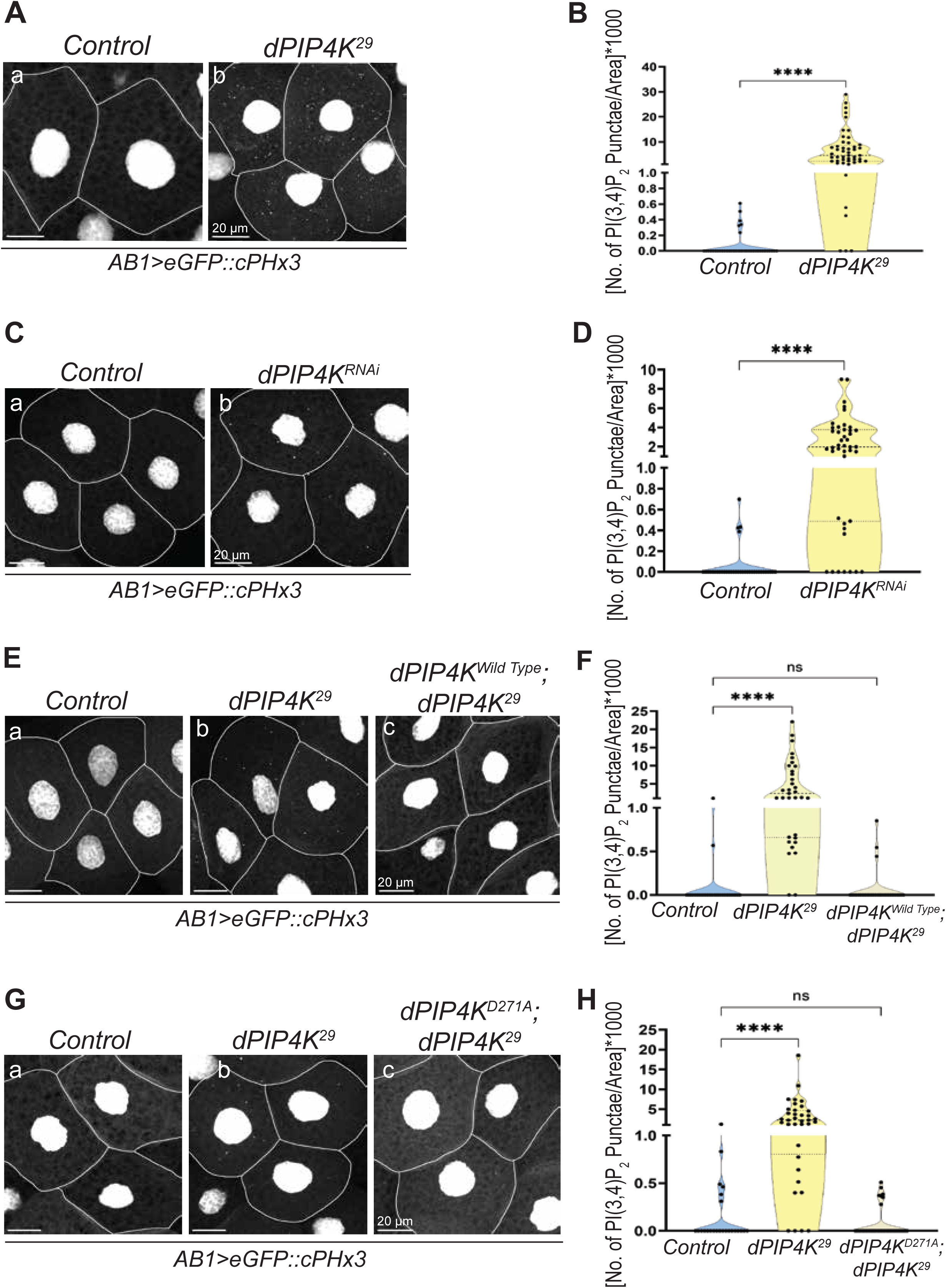
dPIP4K regulates PI(3,4)P_2_ levels in *Drosophila* salivary gland cells. **(A)** Representative confocal z-projections illustrating PI(3,4)P_2_ levels using eGFP::cPHx3 in the salivary glands of wandering third instar larvae from the following genotypes: (a) Control - *AB1>eGFP::cPHx3* and (b) *AB1>eGFP::cPHx3;dPIP4K^29^*. The scale bar is indicated at 20 μm for spatial reference. **(B)** Quantification of PI(3,4)P_2_ levels using eGFP::cPHx3 probe in the salivary glands of wandering third instar larvae from the following genotypes: (a) Control - *AB1>eGFP::cPHx3* (N=12,n=34) and (b) *AB1>eGFP::cPHx3;dPIP4K^29^* (N=13,n=48). Statistical test: Mann Whitney U test (****P value<0.0001). **(C)** Representative confocal z-projections illustrating PI(3,4)P_2_ levels using eGFP::cPHx3 in the salivary glands of wandering third instar larvae from the following genotypes: (a) Control - *AB1>eGFP::cPHx3* and (b) *AB1>eGFP::cPHx3;dPIP4K^RNAi^*. The scale bar is indicated at 20 μm for spatial reference. **(D)** Quantification of PI(3,4)P_2_ levels using eGFP::cPHx3 punctae and in the salivary glands of wandering third instar larvae from the following genotypes: (a) Control - *AB1>eGFP::cPHx3* (N=12,n=36) and (b) *AB1>eGFP::cPHx3;dPIP4K^RNAi^* (N=14,n=43). Statistical test: Mann Whitney U test (****P value<0.0001). **(E)** Representative confocal z-projections illustrating PI(3,4)P_2_ levels using eGFP::cPHx3 in the salivary glands of wandering third instar larvae from the following genotypes: (a) Control - *AB1>eGFP::cPHx3*, (b) *AB1>eGFP::cPHx3;dPIP4K^29^* and (c) *AB1> eGFP::cPHx3,dPIP4K^Wild Type^;dPIP4K^29^.*The scale bar is indicated at 20 μm for spatial reference. **(F)** Quantification of PI(3,4)P_2_ levels using eGFP::cPHx3 punctae and in the salivary glands of wandering third instar larvae from the following genotypes: (a) Control - *AB1>eGFP::cPHx3* (N=9,n=28), (b) *AB1>eGFP::cPHx3;dPIP4K^29^* (N=8,n=33) and (c) *AB1>eGFP::cPHx3; dPIP4K^Wild^ ^Type^,dPIP4K^29^* (N=7,n=29).Statistical test: Kruskal-Wallis Test (****P value < 0.0001 and ns – P value>0.9999). **(G)** Representative confocal z-projections illustrating PI(3,4)P_2_ levels using eGFP::cPHx3 in the salivary glands of wandering third instar larvae from the following genotypes: (a) Control - *AB1>eGFP::cPHx3*, (b) *AB1>eGFP::cPHx3;dPIP4K^29^* and (c) *AB1>eGFP::cPHx3, dPIP4K^D271A^;dPIP4K^29^*.The scale bar is indicated at 20 μm for spatial reference. **(H)** Quantification of PI(3,4)P_2_ levels using eGFP::cPHx3 punctae and in the salivary glands of wandering third instar larvae from the following genotypes: (a) Control - *AB1>eGFP::cPHx3* (N=6,n=24), (b) *AB1>eGFP::cPHx3;dPIP4K^29^* (N=9,n=36) and (c) *AB1> eGFP::cPHx3,dPIP4K^D271A^;dPIP4K^29^* (N=10, n=36). Statistical test: Kruskal-Wallis Test (****P value < 0.0001 and ns – P value>0.9999).

To determine if the observed elevation in PI(3,4)P_2_ levels in *dPIP4K^29^* was specifically due to loss of PIP4K, we reconstituted full length dPIP4K (dPIP4K^Wild^ ^Type^) in *dPIP4K^29^* and observed a full reversal of the phenotype **[Fig 2, E and F]**. Expression levels of the probes were comparable across genotypes **[Supplementary Fig 2E]**. Further, to understand if the observed upregulation in PI(3,4)P_2_ levels required the kinase activity of the enzyme, we reconstituted a kinase dead (non-ATP binding) dPIP4K (dPIP4K^D271A^) in *dPIP4K^29^*. Remarkably and unexpectedly, PI(3,4)P_2_ levels were restored back to those seen in controls **[Fig 2, G and H]** suggesting that regulation of PI(3,4)P_2_ by dPIP4K does not depend on its kinase activity; expression levels of the probes were comparable across genotypes **[Supplementary Fig 2F]**.

### Endomembranous PI(3,4)P_2_ accumulates in *dPIP4K^29^* independent of Class I PI3K activity

*dPIP4K^29^* mutants show elevated PIP_3_ levels on the plasma membrane(Sharma et al., 2019). Therefore, the activity of a 5-phosphatase on PIP_3_ can lead to accumulation of PI(3,4)P_2_ on the plasma membrane, followed by endocytosis onto endomembranes. This hypothesis would predict that PI(3,4)P_2_ on the plasma membrane would be elevated in *dPIP4K^29^*. However, we did not find evidence for such an elevation of PI(3,4)P_2_ at the plasma membrane **[Supplementary Fig 3, D and E]**, suggesting that in *dPIP4K^29^*, plasma membrane PI(3,4)P_2_ might not be the source for elevated PI(3,4)P_2_ at the endomembrane.

To validate this further, we stimulated control salivary glands with 10 μM insulin, a treatment known to increase PIP_3_ in the *Drosophila* salivary glands (Sharma et al., 2019). This should give rise to PI(3,4)P_2_ in a sustained manner but with a temporal delay at the plasma membrane [(Liu et al., 2018);(Goulden et al., 2019)] and if this pool were to be the source for endomembranous pool of PI(3,4)P_2_, one might expect to observe the punctate population of eGFP::cPHx3, with some temporal delay after stimulating glands with insulin (Liu et al., 2018). Using a working concentration of 10 μM insulin, we observed a rapid rise in plasma membrane PIP₃, monitored using the tGPH probe **[Supplementary Fig 3F].** As expected, a time course experiment using the same concentration of insulin on wild type glands led to an increase in PI(3,4)P_2_ at the plasma membrane but there was no corresponding increase of endomembranous PI(3,4)P_2_ levels **[Fig 3A]**. As an alternative approach to test the involvement of PIP_3_ as a precursor for the increased endomembranous PI(3,4)P_2_ in *dPIP4K^29^*, we treated salivary glands with 100 nM wortmannin (Wymann et al., 1996), a concentration at which this compound inhibits Class I PI3K thus reducing PIP_3_ levels at the plasma membrane. If PIP_3_ was indeed the source for the elevated PI(3,4)P_2_ in *dPIP4K^29^*, then pre-treatment with wortmannin is predicted to lead to a reduction in the levels of PI(3,4)P_2._ However, pre-treatment of *dPIP4K^29^* glands with 100 nM of wortmannin did not result in a reduction in PI(3,4)P_2_ punctae **[Fig 3, B and C]**. These findings suggest that the accumulation of PI(3,4)P_2_ in *dPIP4K^29^* is likely independent of Class I PI3K activity.

**Figure 3:**
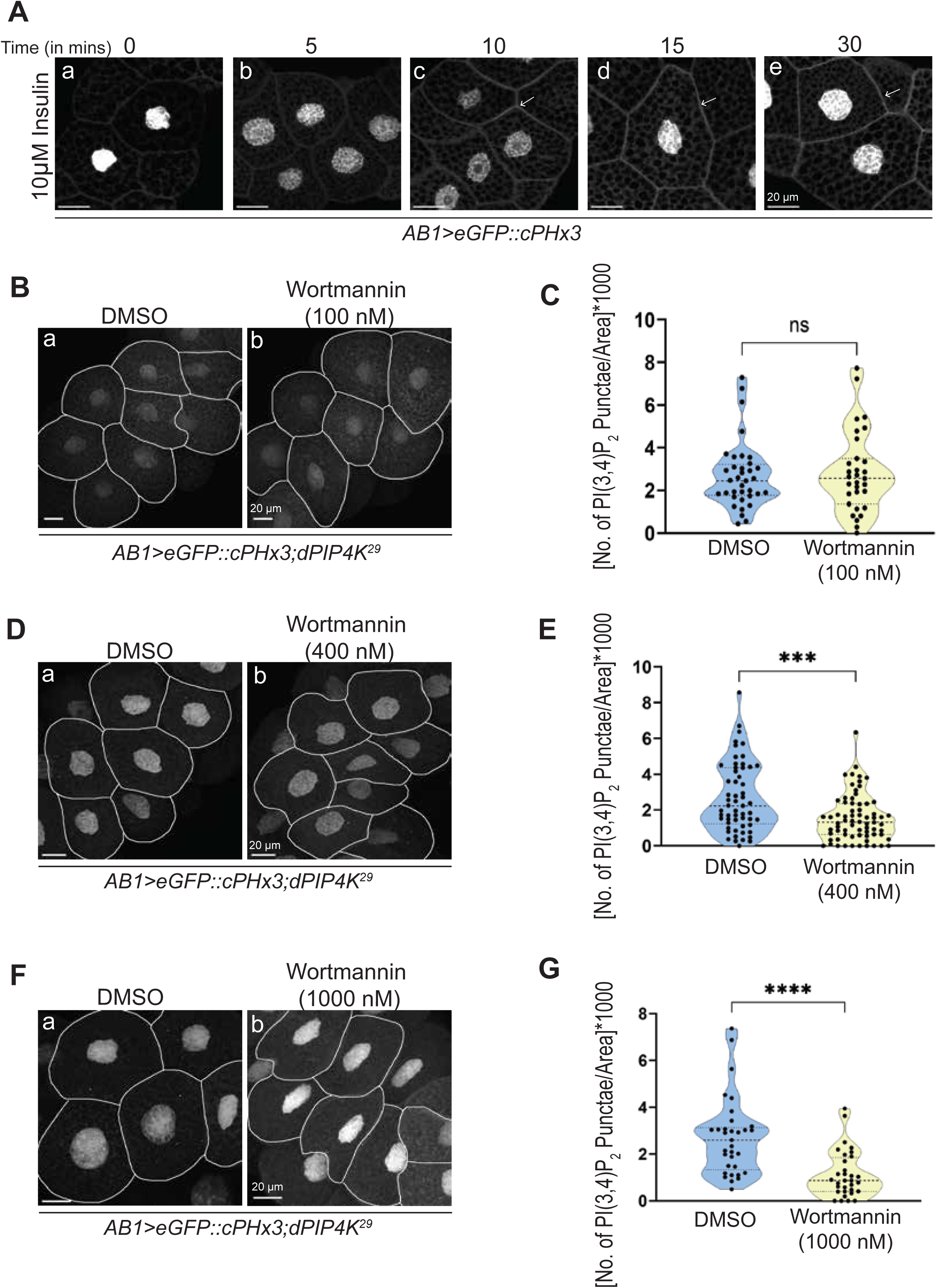
Endomembranous PI(3,4)P_2_ accumulates in *dPIP4K^29^* independent of Class I PI3K activity. **(A)** Representative confocal z-projections illustrating PI(3,4)P_2_ levels using eGFP::cPHx3 in the salivary glands of wandering third instar larvae from the genotype *AB1>eGFP::cPHx3,* insulin stimulated (10 µM) for the following time points - (a) 0 (Unstimulated), (b) 5 mins (c) 10 mins (d) 15 mins and (e) 30 mins. The scale bar is indicated at 20 μm for spatial reference. **(B)** Representative confocal z-projections illustrating PI(3,4)P_2_ levels using eGFP::cPHx3 in the salivary glands of wandering third instar larvae from the following genotypes: (a) *AB1> eGFP::cPHx3;dPIP4K^29^* (DMSO) and (b) *AB1>eGFP::cPHx3;dPIP4K^29^* (Wortmannin-100nM). The scale bar is indicated at 20 μm for spatial reference. **(C)** Quantification of PI(3,4)P_2_ levels using eGFP::cPHx3 punctae and in the salivary glands of wandering third instar larvae from the following genotypes:(a) *AB1>eGFP::cPHx3;dPIP4K^29^* (DMSO) (N=8,n=36) and (b) *AB1>eGFP::cPHx3;dPIP4K^29^* (Wortmannin-100nM) (N=9,n=31). Statistical test: Mann Whitney U test (ns – P value = 0.8563). **(D)** Representative confocal z-projections illustrating PI(3,4)P_2_ levels using eGFP::cPHx3 in the salivary glands of wandering third instar larvae from the following genotypes:(a) *AB1> eGFP::cPHx3;dPIP4K^29^* (DMSO) and (b) *AB1> eGFP::cPHx3;dPIP4K^29^* (Wortmannin-400nM). The scale bar is indicated at 20 μm for spatial reference. **(E)** Quantification of PI(3,4)P_2_ levels using eGFP::cPHx3 punctae and in the salivary glands of wandering third instar larvae from the following genotypes: (a) *AB1>eGFP::cPHx3; dPIP4K^29^*(DMSO) (N=11,n=57) and (b) *AB1>eGFP::cPHx3;dPIP4K^29^* (Wortmannin- 400nM) (N=16,n=70). Statistical test: Mann Whitney U test (***P value<0.001). **(F)** Representative confocal z-projections illustrating PI(3,4)P_2_ levels using eGFP::cPHx3 in the salivary glands of wandering third instar larvae from the following genotypes::(a) *AB1> eGFP::cPHx3;dPIP4K^29^* (DMSO) and (b) *AB1>eGFP::cPHx3;dPIP4K^29^* (Wortmannin-1000nM). The scale bar is indicated at 20 μm for spatial reference. **(G)** Quantification of PI(3,4)P_2_ levels using eGFP::cPHx3 punctae and in the salivary glands of wandering third instar larvae from the following genotypes::(a) *AB1>eGFP::cPHx3;dPIP4K^29^* (DMSO) (N=9,n=33) and (b) *AB1>eGFP::cPHx3;dPIP4K^29^* (Wortmannin- 1000nM) (N=10,n=32). Statistical test: Mann Whitney U test (****P value<0.0001).

### Class II PI3K dependent endomembranous PI(3,4)P_2_ accumulates in *dPIP4K^29^*

During our studies, we noted that pre-treating salivary glands with 400 nM **[Fig 3, D and E]** and 1000 nM **[Fig 3, F and G]** of wortmannin led to a noticeable and dose dependent reduction **[Supplementary Fig 3G]** in PI(3,4)P_2_ punctae. One possible explanation for this differential response (compared to 100nM wortmannin) could be wortmannin’s inhibition profile; at 100 nM, Class I PI3K is completely inhibited and class II is only partially inhibited, whereas at 400 nM and 1000nM, Class I PI3K remains 100% inhibited while Class II PI3K is also inhibited to 50% and 100% respectively (Domin et al., 1997). The *Drosophila* genome has a single gene encoding Class II PI3K (Pi3K68D)(MacDougall et al., 2004). To determine if the PI(3,4)P_2_ punctae in *dPIP4K^29^* are dependent on Class II PI3K activity, we downregulated Pi3K68D using RNAi **[Supplementary Fig 4A]** in *dPIP4K^29^* salivary glands. Under these conditions (*Pi3K68D^RNAi^* in *dPIP4K^29^*), we observed that the elevated PI(3,4)P_2_ punctae in *dPIP4K^29^* were significantly rescued **[Fig 4, A and B]**; the PI(3,4)P_2_ probe was expressed at equal levels in all the genotypes **[Supplementary Fig 4B]**. These findings predict that the elevation of Class II PI3K function in salivary gland cells should increase PI(3,4)P_2_ punctae; i.e., overexpression of Pi3K68D should phenocopy elevated PI(3,4)P_2_ levels in *dPIP4K^29^*. Indeed, we observed elevated levels of endomembranous pool of PI(3,4)P_2_ in glands overexpressing Pi3K68D **[Fig 4, C and D]** with comparable expression levels of the probe across genotypes **[Supplementary Fig 4C]**. The punctae were specific to PI(3,4)P_2_ as a non-binding version of the probe (eGFP::R211LcPHx3) failed to pick up any PI(3,4)P_2_ punctate signal **[Supplementary Fig 4, E and F]**. Expression levels of both the binding and non-binding probes were comparable across genotypes, ruling out probe expression differences as a confounding factor **[Supplementary Fig 4D]**. We next investigated whether loss of dPIP4K alters the levels of PI4P, the substrate for PI(3,4)P₂ synthesis via Class II PI3K activity. Using the mCherry::P4M probe, we found that PI4P-positive punctae were significantly increased in *dPIP4K^29^*salivary glands compared to the controls **[Fig 4, E and F]**. In contrast, PI4P levels at the plasma membrane were significantly reduced **[Supplementary Fig 4H]**. Importantly, expression of the PI4P probe was comparable across all genotypes **[Supplementary Fig 4G]**.

**Figure 4:**
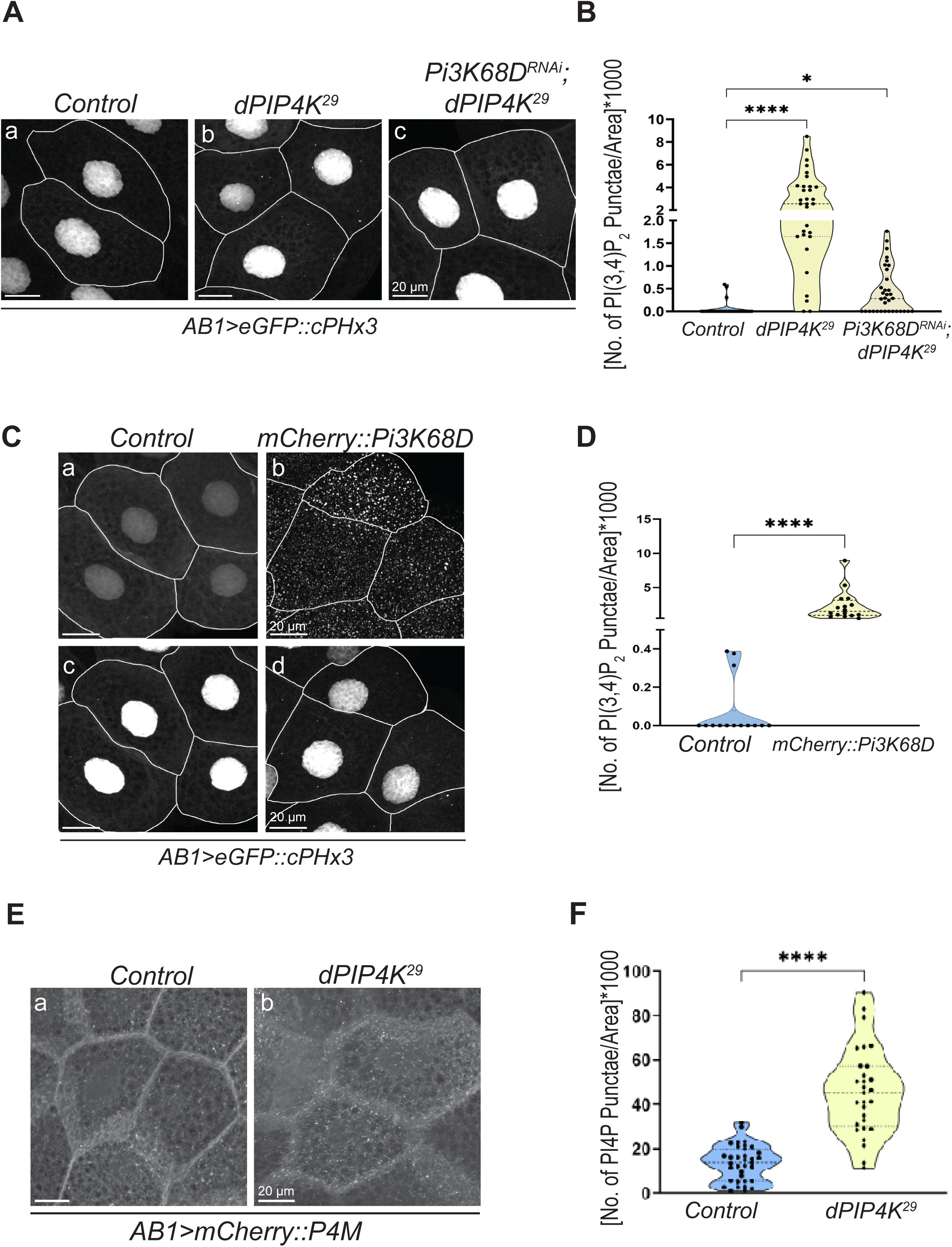
Class II PI3K dependent endomembranous PI(3,4)P_2_ accumulates in *dPIP4K^29^*. **(A)** Representative confocal z-projections illustrating PI(3,4)P_2_ levels using eGFP::cPHx3 in the salivary glands of wandering third instar larvae from the following genotypes: (a) Control - *AB1>eGFP::cPHx3*, (b) *AB1>eGFP::cPHx3;dPIP4K^29^* and (c) *AB1>eGFP::cPHx3,Pi3K68D^RNAi^;dPIP4K^29^*. The scale bar is indicated at 20 μm for spatial reference. **(B)** Quantification of PI(3,4)P_2_ levels using eGFP::cPHx3 punctae and in the salivary glands of wandering third instar larvae from the following genotypes: (a) Control *- AB1> eGFP::cPHx3* (N=11,n=28), (b) *AB1>eGFP::cPHx3;dPIP4K^29^* (N=11,n=31) and (c) *AB1>eGFP::cPHx3,Pi3K68D^RNAi^;dPIP4K^29^*(N=14,n=34). Statistical test: Kruskal-Wallis Test (****P value < 0.0001 and *P value = 0.0210). **(C)** Representative confocal z-projections illustrating PI(3,4)P_2_ levels using eGFP::cPHx3 in the salivary glands of wandering third instar larvae from the following genotypes: (a) and (c) Control - *AB1>eGFP::cPHx3,mCherry*, (b) and (d) *AB1>eGFP::cPHx3,mcherry::Pi3K68D*. The scale bar is indicated at 20 μm for spatial reference. **(D)** Quantification of PI(3,4)P_2_ levels using eGFP::cPHx3 punctae and in the salivary glands of wandering third instar larvae from the following genotypes: (a) and (c) Control-*AB1> eGFP::cPHx3,mCherry* (N=5,n=14), (b) and (d) *AB1>eGFP::cPHx3,mcherry::Pi3K68D* (N=6, n=16). Statistical test: Mann Whitney U test (****P value<0.0001). **(E)** Representative confocal z-projections illustrating PI4P levels using mCherry::P4M in the salivary glands of wandering third instar larvae from the following genotypes: (a) Control - *AB1>mCherry::P4M* and (b) *AB1>mCherry::P4M;dPIP4K^29^*. The scale bar is indicated at 20 μm for spatial reference. **(F)** Quantification of PI4P levels using mCherry::P4M punctae in the salivary glands of wandering third instar larvae from the following genotypes: (a) Control - *AB1>mCherry::P4M* (N=7,n=35) and (b) *AB1>mCherry::P4M;dPIP4K^29^* (N=6,n=29). Statistical test: Unpaired Welch’s t-test (****P value<0.0001)

### Elevated PI(4,5)P_2_ levels give rise to elevated endomembranous levels of PI(3,4)P_2_

One possible mechanism by which PIP4K tunes PI4P levels is by modulating the levels of PI(4,5)P₂ as mentioned in various studies in mammalian systems [(Hinchliffe et al., 2002),(Wang et al.,2019)]. Therefore, we next examined whether elevated PI(4,5)P₂ levels could contribute to the accumulation of endomembranous PI4P and PI(3,4)P₂. To quantify PI(4,5)P₂ levels, we utilized the PH-PLCδ::mCherry biosensor. We observed a significant increase in PI(4,5)P₂-positive punctae in dPIP4K RNAi salivary glands **[Fig 5, A and B],** while plasma membrane-associated PI(4,5)P₂ levels decreased **[Supplementary Fig 5B].** Probe expression was comparable across genotypes **[Supplementary Fig 5A].**

**Figure 5:**
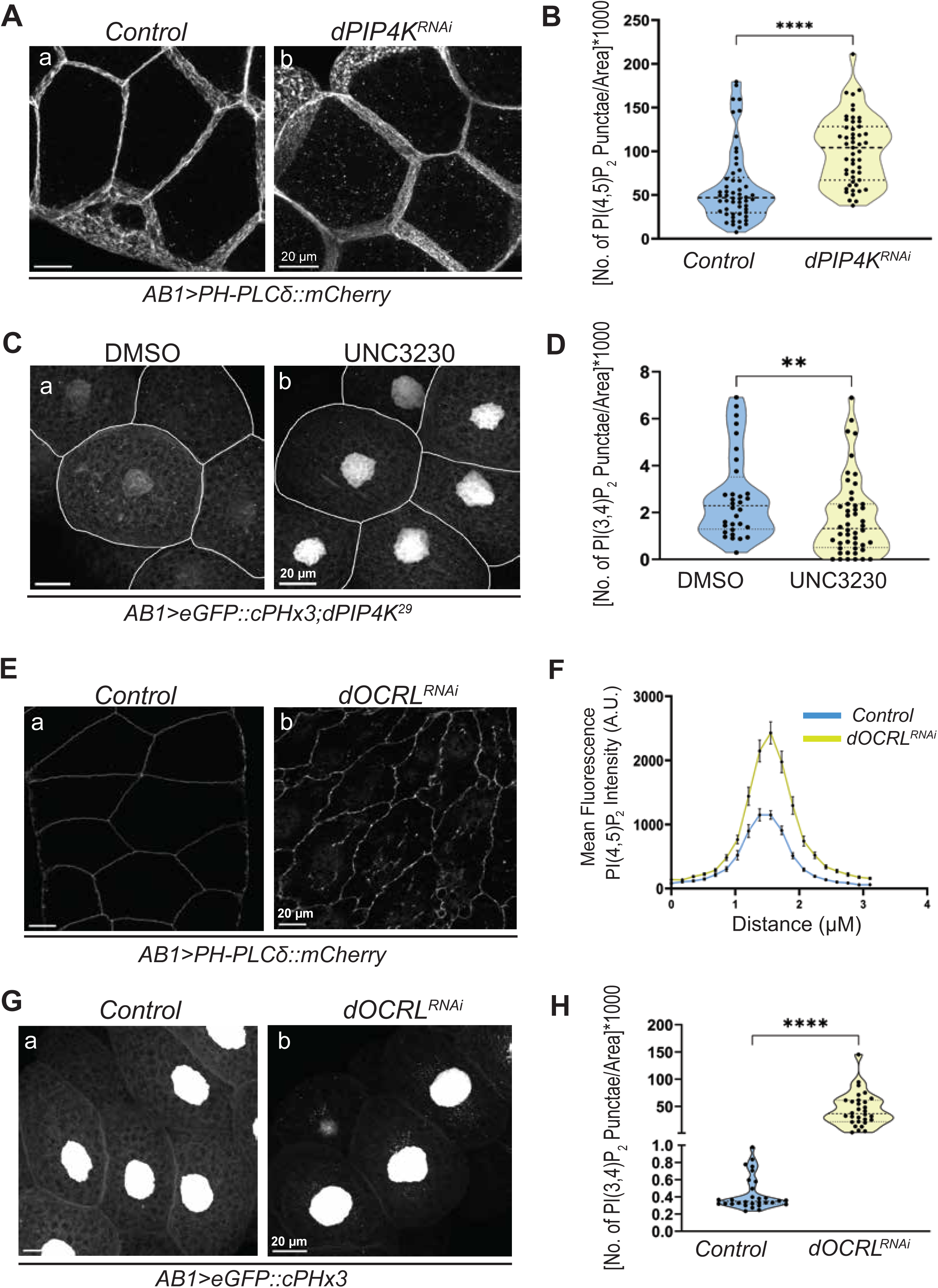
Elevated PI(4,5)P_2_ levels give rise to elevated endomembranous levels of PI(3,4)P_2_. **(A)** Representative confocal z-projections illustrating PI(4,5)P_2_ levels using PH-PLCδ::mCherry in the salivary glands of wandering third instar larvae from the following genotypes: (a) Control - *AB1>PH-PLCδ::mCherry* and (b) *AB1>PH-PLCδ::mCherry;dPIP4K^RNAi^*.The scale bar is indicated at 20 μm for spatial reference. **(B)** Quantification of PI(4,5)P_2_ levels using PH-PLCδ::mCherry punctae and in the salivary glands of wandering third instar larvae from the following genotypes: (a) Control - *AB1>PH-PLCδ::mCherry* (N=11,n=54) and (b) *AB1> PH-PLCδ::mCherry;dPIP4K^RNAi^* (N=12,n=51). Statistical test: Mann- Whitney U test (****P value<0.0001). **(C)** Representative confocal z-projections illustrating PI(3,4)P_2_ levels using eGFP::cPHx3 in the salivary glands of wandering third instar larvae from the following genotypes: (a) *AB1> eGFP::cPHx3;dPIP4K^29^*(DMSO) and (b) *AB1>eGFP::cPHx3;dPIP4K^29^*(UNC3230 −10 µM). The scale bar is indicated at 20 μm for spatial reference. **(D)** Quantification of PI(3,4)P_2_ levels using eGFP::cPHx3 punctae and in the salivary glands of wandering third instar larvae from the following genotypes: (a) *AB1>eGFP::cPHx3; dPIP4K^29^*(DMSO) (N=11,n=32) and (b) *AB1>eGFP::cPHx3;dPIP4K^29^* (UNC3230 −10 µM) (N=12,n=48). Statistical test: Mann Whitney U test (**P value=0.0080). **(E)** Representative confocal z-projections illustrating PI(4,5)P_2_ levels using PH-PLCδ::mCherry in the salivary glands of wandering third instar larvae from the following genotypes: (a) Control - *AB1>PH-PLCδ::mCherry* and (b) *AB1>PH-PLCδ::mCherry;OCRL^RNAi^*. The scale bar is indicated at 20 μm for spatial reference. **(F)** Quantification of PI(4,5)P_2_ levels using PH-PLCδ::mCherry in the salivary glands of wandering third instar larvae from the following genotypes: (a) Control - *AB1>PH-PLCδ::mCherry* (N=4, n=20) and (b*) AB1>PH-PLCδ::mCherry;OCRL^RNAi^* (N=4, n=20). Mean ± SEM is plotted from 4 different salivary glands with 5 cells each. The x-axis represents the distance (µm) along the ROI drawn and the y-axis indicates the mean fluorescence PI(4,5)P_2_ intensity (arbitrary units, AU). **(G)** Representative confocal z-projections illustrating PI(3,4)P_2_ levels using eGFP::cPHx3 in the salivary glands of wandering third instar larvae from the following genotypes: (a) Control - *AB1>eGFP::cPHx3* and (b) *AB1>eGFP::cPHx3;OCRL^RNAi^*. The scale bar is indicated at 20 μm for spatial reference. **(H)** Quantification of PI(3,4)P_2_ levels using eGFP::cPHx3 punctae and in the salivary glands of wandering third instar larvae from the following genotypes: (a) Control - *AB1>eGFP::cPHx3* (N=9,n=31) and (b) *AB1> eGFP::cPHx3;OCRL^RNAi^* (N=9,n=31). Statistical test: Mann Whitney U test (****P value<0.0001).

As an additional test of this hypothesis, we inhibited PIP5K activity using UNC3230, a potent chemical inhibitor of PIP5K. Treatment of wild-type *Drosophila* salivary glands with 10 μM UNC3230 for 30 minutes resulted in a marked reduction in plasma membrane localization of the PI(4,5)P₂ sensor PH-PLCδ::mCherry, confirming effective depletion of PI(4,5)P₂ [Supplementary Fig 5, C and D]. Under identical treatment conditions, dPIP4K mutant salivary glands exhibited a significant reduction in PI(3,4)P₂-positive punctae, as measured using the eGFP::cPHx3 probe [**Fig 5, C and D**]. These observations suggest that elevated PI(4,5)P₂ levels in dPIP4K depleted cells contributes to the accumulation of endomembranous PI(3,4)P₂.

Our model predicts that increasing PI(4,5)P₂ levels independent of dPIP4K depletion should similarly elevate PI(3,4)P₂ punctae. To test this, we downregulated dOCRL, a 5-phosphatase that degrades PI(4,5)P₂ [Supplementary Fig 5E]. As expected, dOCRL depletion resulted in elevated PI(4,5)P₂ levels at both the plasma membrane and within the cytoplasm [**Fig 5, E and F**], while probe expression remained unchanged [Supplementary Fig 5F]. Strikingly, dOCRL knockdown also caused a dramatic increase in the endomembranous pool of PI(3,4)P₂ [**Fig 5, G and H**], despite equivalent probe expression between control and RNAi glands [Supplementary Fig 5G]. In contrast, plasma membrane-associated PI(3,4)P₂ levels were unaffected by dOCRL depletion [Supplementary Fig 5H]. Together, these findings indicate that the endomembranous pool of PI(3,4)P₂ is generated from an endomembranous pool of PI(4,5)P₂ and support a model in which elevated PI(4,5)P₂ drives the accumulation of intracellular PI(3,4)P₂.

## Discussion

The generation of plasma membrane PI(3,4)P₂ through the dephosphorylation of PIP_3_ generated by Class I PI3K activity is well established as is its role in recruiting effector proteins that drive actin remodelling (Montaño-Rendón et al., 2022) and cell migration (Feng & Yu, 2021). By contrast far less is understood about the source of endomembrane PI(3,4)P₂ pools owing to its low concentration, rapid turnover and challenges in detection. A previous study had described the localization of a TAPP1 based probe to endocytic structures (He et al., 2017). However, Goulden et al., suggested that this localisation of the probe to endocytic structures was due to the presence of a clathrin-binding motif in the C terminus end of the TAPP1 probe(Goulden et al., 2019). In this study, we report the development and use *in vivo* of a high avidity probe eGFP::cPHx3 (originally described by Goulden et.al) in *Drosophila*; this probe, based on the C-terminal PH domain of TAPP1 does not have a clathrin binding motif. However, we unequivocally find the presence of an endomembranous pool of PI(3,4)P_2_ using this probe. This finding strongly suggests that in addition to plasma membrane pool, *Drosophila* cells also contain an endomembranous pool of PI(3,4)P_2_. This endomembranous pool of PI(3,4)P_2_ is not readily detected in cells under resting conditions. However, in this study we noted that stimulation of cultured *Drosophila* S2R+ cells with H_2_O_2_ resulted in an increase in the endomembranous pool of PI(3,4)P_2_. These findings suggest that the endomembrane pool of PI(3,4)P_2_ may serve as a signalling lipid during response to stimuli.

What might be the signalling function of an endomembrane pool of PI(3,4)P_2_Recent studies strongly suggest that the spatial and temporal dynamics of PI(3,4)P₂ signaling are critical determinants of its function. At the plasma membrane, PI(3,4)P₂ promotes cell growth and survival through the recruitment of PH-domain–containing effectors such as TAPP1 and TAPP2, consistent with studies showing enhanced AKT activation in TAPP mutant mice (Landego et al., 2012). In addition, Liu at al., have demonstrated a new independent signaling function of PI(3,4)P₂; i.e., isoform- and site-specific membrane recruitment and activation of Akt2. They propose that PIP_3_ is a transient signal confined at the plasma membrane, whereas PI(3,4)P₂ is a more sustained signal that operates at both the plasma membrane and early endosomes (Liu et al., 2018).

What are the biochemical regulators of endomembranous pool of PI(3,4)P_2_? Our study identified dPIP4K as a novel regulator of PI(3,4)P₂ in *Drosophila*; loss of dPIP4K results in the accumulation of PI(3,4)P₂ on endomembranes. Although PIP4K has been reported to phosphorylate PI3P and generate PI(3,4)P_2_, this enzymatic reaction cannot explain the elevation of PI(3,4)P_2_ levels seen in cells depleted of the enzyme. Our finding that the elevated PI(3,4)P_2_ levels in dPIP4K depleted cells could be rescued by reconstitution with a kinase dead dPIP4K transgene implies that the mechanism through which the enzyme controls PI(3,4)P_2_ levels is non-enzymatic. Multiple studies have shown that depletion of PIP4K leads to an elevation of PIP_3_ [(D. G. Wang et al., 2019);(Sharma et al., 2019)] raising the possibility that the elevated PI(3,4)P_2_ may rise by dephosphorylation of PIP_3_. Our findings in this study indicate that this is unlikely to be the case (i) elevation of PIP_3_ levels at the plasma membrane by insulin stimulation did not lead to a concomitant elevation of PI(3,4)P_2_ at the endomembrane. This observation is consistent with Goulden et al., (Goulden et al., 2019) who failed to detect endosomal PI(3,4)P_2_ localization in serum-stimulated 293A cells or after insulin stimulation in HeLa cells. (ii) Selective inhibition of Class I PI3K by low nanomolar concentrations of wortmannin failed to decrease the elevated PI(3,4)P_2_ levels in PIP4K depleted cells.

In this study, we found evidence of a key role for Class II PI3K in regulating endomembrane PI(3,4)P_2_ in salivary gland cells. (i) Overexpression of Class II PI3K increased PI(3,4)P_2_ puncta Treatment with pharmacologically relevant concentrations of wortmannin that can inhibit Class II PI3K could reverse PI(3,4)P_2_ puncta accumulated in PIP4K depleted cells (iii) RNAi depletion of Class II PI3K in PIP4K depleted cells could reverse PI(3,4)P_2_ levels. Together, these findings suggest that increased Class II PI3K activity in dPIP4K depleted cells lead to elevated PI(3,4)P_2_ levels.

A central question arising from this study is how PIP4K regulates Class II PI3K dependent PI(3,4)P₂ production at endomembranes. Our findings provide evidence that altered endomembranous PI(4,5)P_2_ levels contribute to this regulation **[Fig 6]**. (i) Loss of dPIP4K resulted in elevated endomembranous pools of PI(4,5)P₂, (ii) pharmacological depletion of PI(4,5)P₂ reduced the accumulation of PI(3,4)P₂ in dPIP4K mutant glands and, (iii) increasing PI(4,5)P₂ levels through depletion of the 5-phosphatase dOCRL was sufficient to elevate endomembranous PI(3,4)P₂.

**Figure 6:**
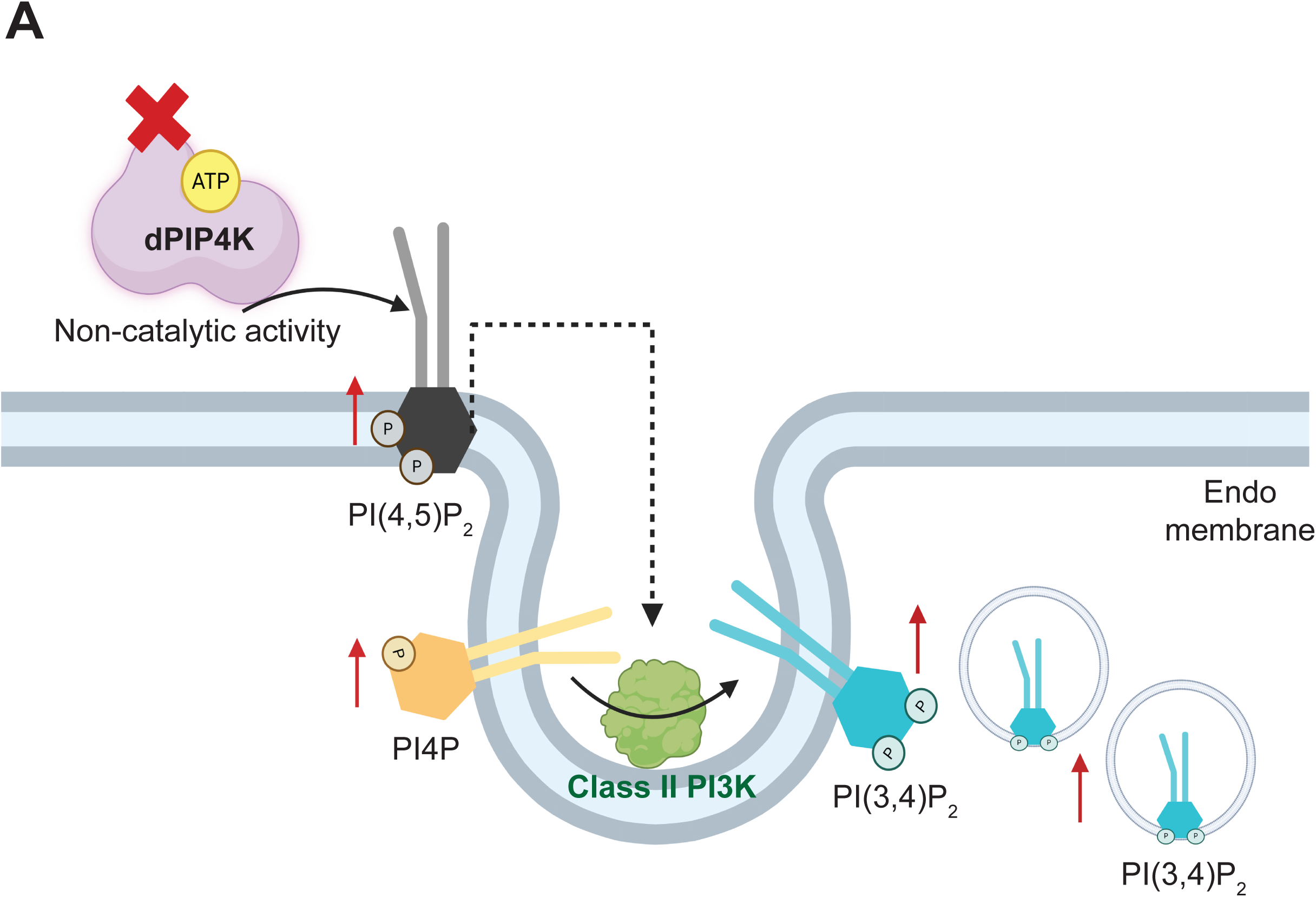
Schematic depicting the non-catalytic regulation of PI(3,4)P₂ levels by dPIP4K on an endomembrane. **(A)** In an otherwise wild type background, dPIP4K regulates PI(4,5)P₂ levels through a catalytically independent mechanism. In a *dPIP4K* null mutant background (denoted by red cross), this mode of regulation is lost and the levels of PI(4,5)P₂ are elevated. This increase in PI(4,5)P₂ may subsequently elevate PI4P levels and enhance class II PI3K–mediated conversion of PI4P to PI(3,4)P₂. As a result, there is an increased pool of endomembranous PI(3,4)P₂. However, it remains unclear how PI(4,5)P₂ is converted into PI4P in this context. Additionally, the elevated PI(4,5)P₂ may also allosterically activate class II PI3K (denoted by black dotted arrow), further promoting the conversion of PI4P into PI(3,4)P₂.

Elevated PI(4,5)P₂ may influence PI(3,4)P₂ production through multiple mechanisms. In addition to the increase in PI4P observed in our study, which could enhance PI(3,4)P₂ synthesis by increasing substrate availability for Class II PI3K, elevated PI(4,5)P₂ may also directly regulate PI3K-C2α activity. A recent study proposed an autoregulatory mechanism for PI3K-C2α involving coincidence detection at endocytic sites (H. Wang et al., 2018). In the cytosol, PI3K-C2α is maintained in an inactive state by folding of the PX-C2 module back onto its kinase domain. Upon membrane recruitment, the N-terminal region of PI3K-C2α interacts with assembled clathrin [(Gaidarov et al., 2001);(Posor et al., 2013);(Domin et al., 2000)] and the C-terminal PX-C2 domain now binds to PI(4,5)P_2_. This conformational change relieves autoinhibition and allows the substrate to access the catalytic site, enabling localized PI(3,4)P_2_ synthesis (H. Wang et al., 2018). Distinguishing the relative contributions of PI4P substrate availability and PI(4,5)P₂-dependent regulation of PI3K-C2α will be important for fully understanding how PIP4K controls Class II PI3K signalling and PI(3,4)P₂ homeostasis at endomembranes.

While the mechanism of action and the regulators of Class I PI3K has been extensively studied, the regulators of Class II PI3K has remained elusive. In our study, we identified a potential novel regulator of Class II PI3K activity that is extrinsic to the enzyme, suggesting an additional layer of control. Recent analysis has shown that PIP4K is a metazoan specific enzyme (Krishnan et al., 2025). Likewise, Class II PI3K is also a metazoan specific lipid kinase [as reviewed in (Ray et al., 2024)] leading to the possibility of an evolutionary link between the two to regulate phosphoinositide signalling in metazoans.

Looking forward, our findings open new directions for studying PI(3,4)P₂ signalling at endomembranes. Future work will be needed to determine whether PIP4K similarly regulates PI(3,4)P₂ in other cell types and physiological contexts. It will be crucial to explore the downstream consequences of heightened PI(3,4)P₂ at endo-membranes in terms of the effector proteins it binds to as well as signalling pathways it maps onto. Given the central role of PI(3,4)P₂ in both growth-promoting and growth-restraining pathways, uncovering how PIP4K maintains its balance is likely to have broad implications for cell signalling, metabolism, and disease.

## Materials and Methods

### Fly husbandry

*Drosophila melanogaster* was grown in a constant temperature laboratory incubator (25°C) on a standard fly media, the composition of which is mentioned in a table below.

**Table 1:** List of ingredients used for the preparation of standard fly media.

| Ingredients | Amount |
| --- | --- |
| 1. Corn flour | 80 g |
| 2. D-Glucose | 20 g |
| 3. Sugar | 40 g |
| 4. Agar | 08 g |
| 5. Yeast powder | 15 g |
| 6. Propionic acid | 4 ml |
| 7. TEGO<br>(Methylparahydroxy benzoate) | 0.7 g |
| 8. Orthophosphoric acid | 0.6 ml |

Experimental fly crosses were set at 25°C in vials under non-crowding conditions with 50% relative humidity under no internal illumination in the incubator. The *Drosophila* strains used in the experiment is listed below – *w^1118^* (Wild type strain), *dPIP4K^29^* (homozygous null mutant of dPIP4K generated in Raghu lab), *UAS-eGFP::cPHx3/Tm6Tb* (Lab generated), *UAS-eGFP::R211LcPHx3/Tm6Tb* (Lab generated), *Act5C-Gal4/CyOGFP* (BL 3953)*, AB1Gal4* (BDSC 1824), *UAS-dPlP4K^RNAi^* (BDSC 65891),*UAS-dPIP4K^WildType^*(Lab generated), *UAS-dPIP4K^D271A^*(Lab generated),*UAS-mCherry,UAS-Pi3K68D^RNAi^*(kind gift from Amy Kiger, UCSD) and *UAS-mCherry::Pi3K68D^2.1^/CyO* (kind gift from Amy Kiger, UCSD)*, UAS-dOCRL^RNAi^*(VDRC, KK110796), *UAS-4cPH-PLCδ1::mCherry* (Patrick Verstreken, VIB, Belgium) and *UAS-mCherry::P4M/Tm6Tb* (Available in the lab).

All experimental procedures, including animal care and use, were conducted under the approval of the Institutional Biosafety Committee (IBSC). Experimental designs incorporated *Drosophila* of both sexes to minimize potential sex-based variation and eliminate sex bias. The Gal4–UAS system (Brand & Perrimon, 1993) was employed to achieve spatially and temporally controlled expression of the transgene.

### DNA constructs and transgenic fly generation

The eGFP::cPHx3 sequence was obtained from Goulden et al.,2019 (Goulden et al., 2019). The sequence for eGFP::R211LcPHx3 was obtained by changing the codon encoding arginine (R) to leucine (L) in all the three tandem repeats. Both the sequences were then provided to GenScript (RRID: SCR_002891) for custom gene/DNA synthesis followed by subsequent incorporation into a *Drosophila* expression vector pUAST-attB (RRID: DGRC_1419). Transgenic fly lines were generated using phiC31 integrase-mediated transgenesis in either attP40 (BDSC 25709) or attP2 (BDSC 25710) flies.

### S2R+ cells: culturing and transfection

*Drosophila* S2R+ cells (RRID: CVCL_Z831) stably expressing Actin-Gal4 were maintained in Schneider’s insect medium (SIM) (HiMedia, Cat# IML003A) supplemented with 10% of a non-heat inactivated fetal bovine serum (FBS, Gibco, Cat# 16000044) and in the presence of glutamine and antibiotics such as penicillin and streptomycin (1:100). Effectene (Qiagen, Cat# 301425) was used to transfect 0.5m cells (for both imaging and western) according to the manufacturer’s instructions. After 36 hours of transfection, cells were fixed with 2.5% paraformaldehyde (PFA) (Electron Microscopy Sciences, Cat# 15710) and imaged to observe for GFP fluorescence using a 60X 1.4 NA objective in Olympus Confocal Laser Scanning Microscope Fluoview FV3000 (RRID:SCR_017015).

### Starvation treatment and H_2_O_2_ treatment in S2R+ cells

Following transfection with the eGFP::cPHx3 and eGFP::R211LcPHx3 probe, cells were dislodged and plated onto confocal dishes and allowed to settle for 2 hours at 25°C in 200 ul of fresh SCM. Post the attachment of cells to the dish surface, the existing media was taken out and the desired treatment was given.

- **For starvation** - Control cells were kept in SCM for 30 mins and treated cells were kept in PBS for 30 mins.
- **For H_2_0_2_** - Control cells were kept in SCM for 30 mins and treated cells were kept in H_2_0_2_ (10 mM) in SCM for 30 mins.

Post the desired treatment, cells were immediately fixed with 2.5% paraformaldehyde (PFA) (Electron Microscopy Sciences, Cat# 15710) and imaged to observe for GFP fluorescence using a 60X 1.4 NA objective in Olympus Confocal Laser Scanning Microscope Fluoview FV3000 (RRID:SCR_017015).

### qPCR

Total RNA was isolated from 5 *Drosophila* 3^rd^ instar wandering larvae using the TRIzol reagent (Life Technologies, Cat# 15596018). 1000ng of the extracted RNA was then treated with DNase I (Amplification grade; Thermo Fisher Scientific, Cat# 18068015) to eliminate any genomic DNA contamination. First-strand cDNA synthesis was carried out using SuperScript II Reverse Transcriptase (Thermo Fisher Scientific, Cat# 18064014) along with random hexamer primers (Thermo Fisher Scientific, Cat# N8080127). Primers targeting exon–exon junctions were designed based on standard qPCR design guidelines. Quantitative PCR (qPCR) was performed using cDNA samples and primers using Power SYBR Green PCR Master Mix (Applied Biosystems, Cat# 4367659) on an Applied Biosystems 7500 Fast Real-Time PCR System (RRID:SCR_018051). Expression levels were normalized to the ribosomal protein 49 (RP49) a housekeeping gene, thus serving as the internal reference control. Additionally, a no reverse transcription control and a non-template control was also included for each genotype to rule out amplification coming from genomic DNA or non-specific contaminants.

### Protein sample preparation

- **Salivary glands** - 6 stage-matched *Drosophila* larvae were dissected for salivary glands and the samples were sonicated in larval lysis buffer (3 µl/gland) with freshly added PIC (Protease Inhibitor Cocktail) (Roche, Cat# 4693116001) and PS (PhosStop) (Roche, Cat# 4906845001), the detailed composition of lysis buffer is mentioned in Swarna et al., 2019 (Mathre et al., 2019). Following lysis, the samples were boiled at 95°C for 10 min.
- **S2R+ cells** - Lysates of S2R+ cells post transfection expressing the desired construct were prepared by pelleting the cells at 1000 rpm/4°C/15 mins. The pellet was subsequently washed using ice cold 1X PBS (2 times). Following this, the cell pellet was lysed in the Lamelli buffer, heated at 95°C for 10 min.

### Western Blotting

Protein extracts were separated using an SDS-PAGE and electro blotted onto a nitrocellulose filter membrane [Hybond-C Extra; (GE Healthcare)] using a wet transfer apparatus (Bio-Rad). The membrane was blocked using 10% Blotto (Santa Cruz Biotechnology, Cat# sc-2325) in phosphate buffer saline (PBS) with 0.1%Tween 20 (Sigma-Aldrich, Cat# P1379) (0.1% PBST) for an hour at room temperature (RT) on an orbital shaker. Primary antibody incubation was done overnight at 4°C using appropriate antibody dilutions as listed in the table 2. Following this, the membrane was washed in 0.1% PBST (3 times) at RT and incubated with 1: 10,000 dilutions of appropriate secondary antibody (Jackson Immuno Research Laboratories) coupled to horseradish peroxidase (HRP) (Donkey anti rabbit, Cat # 711-035-152, RRID: AB_10015282), (Goat anti mouse, Cat# 115-035-003, RRID: AB_10015289) at RT for 2 h on an orbital shaker. The membranes were washed three times with PBST, developed using Clarity western ECL substrate (Bio-Rad Cat# 1705061), and imaged with the LAS 4000 ImageQuant system (GE Healthcare; RRID:SCR_014246).

**Table 2:**
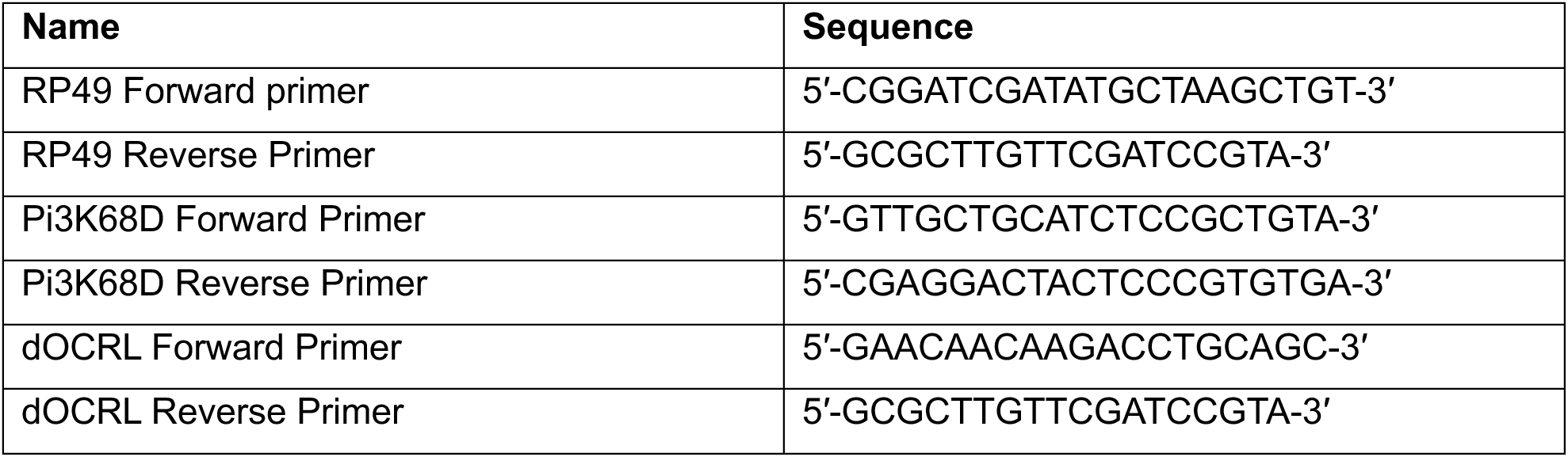
List of primers used for qPCR measurement of Pi3K68D and dOCRL.

**Table 3:** List of primary antibodies used for western blotting.

| **Primary Antibody** | **Dilution** | **Catalogue Number** | **RRID** | **Source** |
| --- | --- | --- | --- | --- |
| Rabbit polyclonal anti-α-Actin | 1:1000 | A5060 | RRID: AB_476738 | Sigma-Aldrich |
| Mouse polyclonal anti-β-Tubulin | 1:4000 | E7-c | RRID: AB_528499 | Developmental Studies Hybridoma Bank (DSHB) |
| Rabbit polyclonal anti-dPIP4K | 1:1000 | - | - | Generated for the laboratory by NeoBiolab |
| Mouse polyclonal anti-GFP | 1:2000 | sc-9996 | RRID: AB_627695 | Santa Cruz Biotechnology |

### Treatments in salivary glands

- **For Insulin time course** - Wandering third instar larvae were dissected one at a time and glands were immediately dropped in one well of a 4-well plate containing either only PBS or PBS+1µM insulin (Bovine Insulin, Sigma, Cat# I6634) and incubated for multiple time points such as 5, 10, 15 and 30 minutes at 25°C following which the glands were transferred to a well containing 4% PFA in PBS to fix for 15 minutes at room temperature and then washed with 1X PBS (2 times). Finally, the glands were mounted in 70% glycerol in PBS and imaged to observe for GFP fluorescence using a 60X 1.4 NA objective in Olympus Confocal Laser Scanning Microscope Fluoview FV3000 (RRID:SCR_017015).
- **For Wortmannin treatment** - Wandering third instar larvae were dissected and glands were immediately dropped in one well of a 4-well plate containing either DMSO in Schneider’s insect media or Wortmannin (Sigma-Aldrich, Cat# W1628) at desired concentrations (100nM, 400nM or 1000nM) in Schneider’s insect media and incubated for 10 minutes at room temperature following which the glands were transferred to a well containing 4% PFA in PBS and fixed for 15 minutes at room temperature and then transferred to wells containing only 1X PBS for washes (2 times). Finally, the glands were mounted in 70% glycerol in PBS and imaged to observe for GFP fluorescence using a 60X 1.4 NA objective in Olympus Confocal Laser Scanning Microscope Fluoview FV3000 (RRID:SCR_017015).
- **For UNC3230 treatment** - Wandering third instar larvae were dissected and glands were immediately dropped in one well of a 4-well plate containing either DMSO in Schneider’s insect media or UNC-3230 (TOCRIS, Cat# 5271) at the desired concentration (10 µM) in Schneider’s insect media and incubated for 30 minutes at room temperature. Following this, 16% Paraformaldehyde (PFA) was added directly to the treatment well such that the final concentration of PFA is 4%, and glands were incubated in this for 20 minutes at room temperature. The fixed glands were washed once in 1X PBS for 10 minutes and fat bodies attached to glands were cleaned prior to mounting. Finally, the glands were mounted in 70% glycerol in PBS and imaged to observe for GFP fluorescence using a 60X 1.4 NA objective in Olympus Confocal Laser Scanning Microscope Fluoview FV3000 (RRID:SCR_017015).

### Analysis

#### • Punctae quantification

The 3D images were stacked to give one 2D image using ZProject in ImageJ. The 2D images were then analysed for the number of punctae either counted manually or by using the 3D object counter plugin in ImageJ. The number of punctae were normalised to the area of the cell. The value thus obtained was multiplied by 1000 and plotted for the respective genotypes.

#### • Plasma membrane to Cytoplasm mean fluorescence intensity (MFI) Quantification

Confocal 3D slices were manually curated to generate maximum z-projections of middle few planes (2-3 slices) of cells. Thereafter, line profiles were drawn across clearly identifiable plasma membrane regions and their adjacent cytosolic regions and ratios of mean intensities for these line profiles were calculated for each cell. For salivary glands, about 5-6 cells from multiple glands were analysed.

### Sampling and Statistical Analysis

Each experiment was performed at least twice with multiple biological replicates. All statistical analyses were conducted using GraphPad Prism (RRID:SCR_002798). Data normality or log-normality was assessed using the Shapiro–Wilk test (*p* > 0.05). For comparisons between two groups, the Mann–Whitney *U* test was applied if the data did not follow a normal distribution, whereas an unpaired *t*-test with Welch’s correction was used for normally distributed data. When experiments involved more than two biological groups and the data were non-normal, the Kruskal–Wallis test was employed.

## Acknowledgements

This work was supported by the Department of Atomic Energy, Government of India, under Project Identification No. RTI 4006. We thank the Imaging, DNA sequencing and *Drosophila* facility at NCBS for support. We thank Amy Kiger (UCSD) for her generous support in providing the Class II PI3K RNAi and overexpression lines used in this study. We also thank the members of the Padinjat laboratory especially Avishek Ghosh, Sanjeev Sharma and Swarna Mathre for help, line generation and advice during this study.

