## Supplementary Figures 1-5 for "Phosphatidylinositol 5 phosphate 4 kinase regulates phosphatidylinositol 3,4 bisphosphate levels *in vivo*"

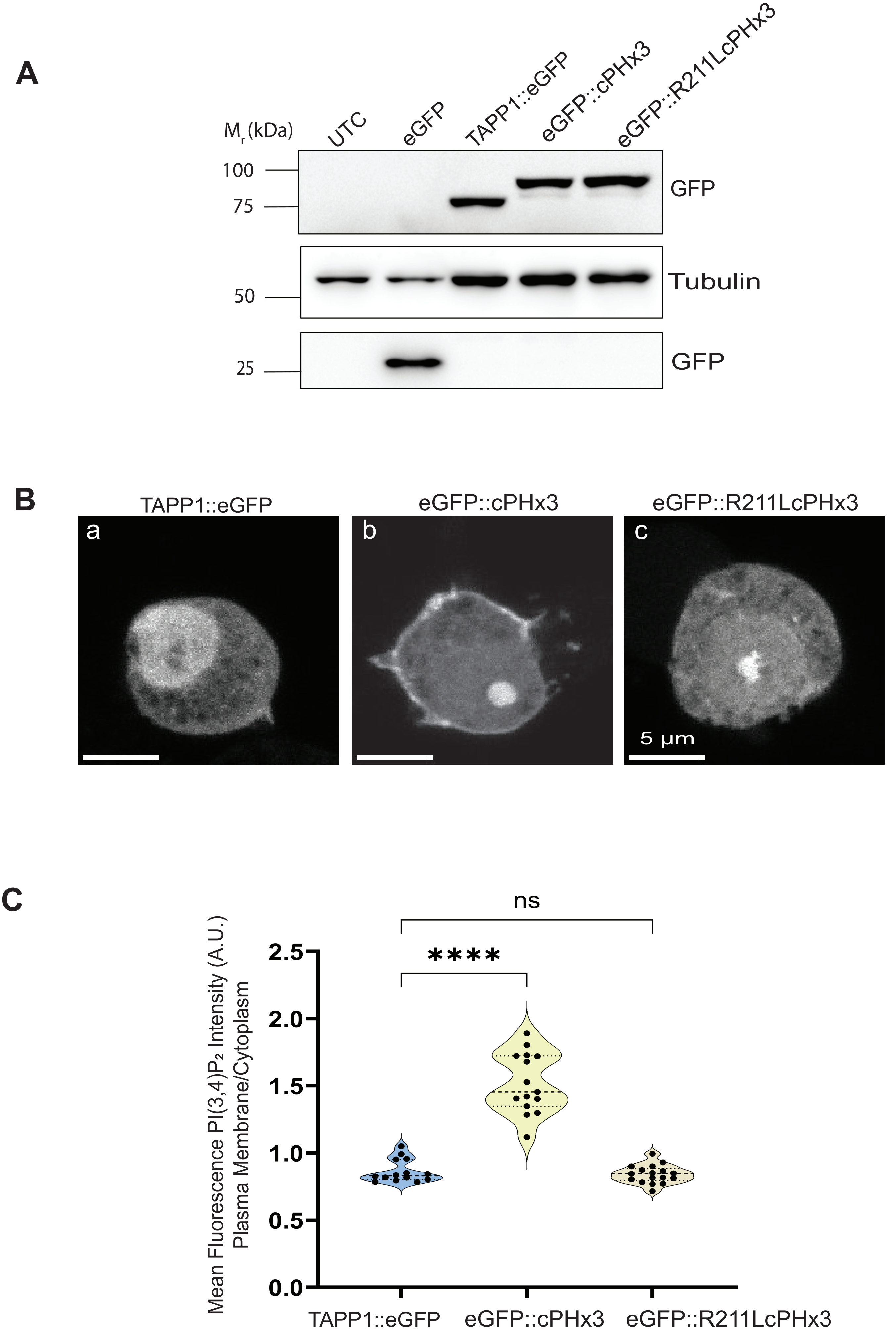


*Venugopal et al.,Supplementary Figure 1*

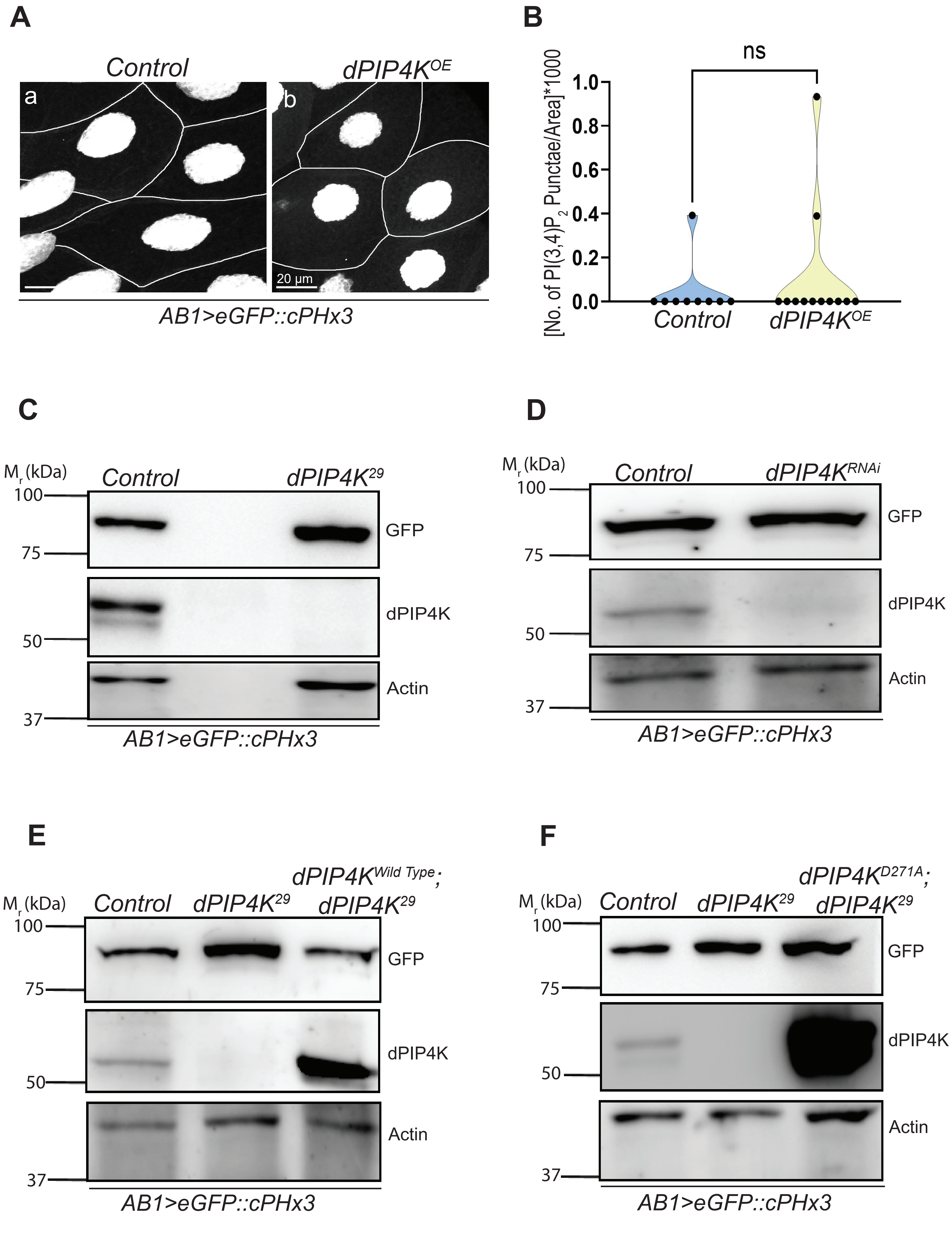


*Venugopal et al.,Supplementary Figure 2*

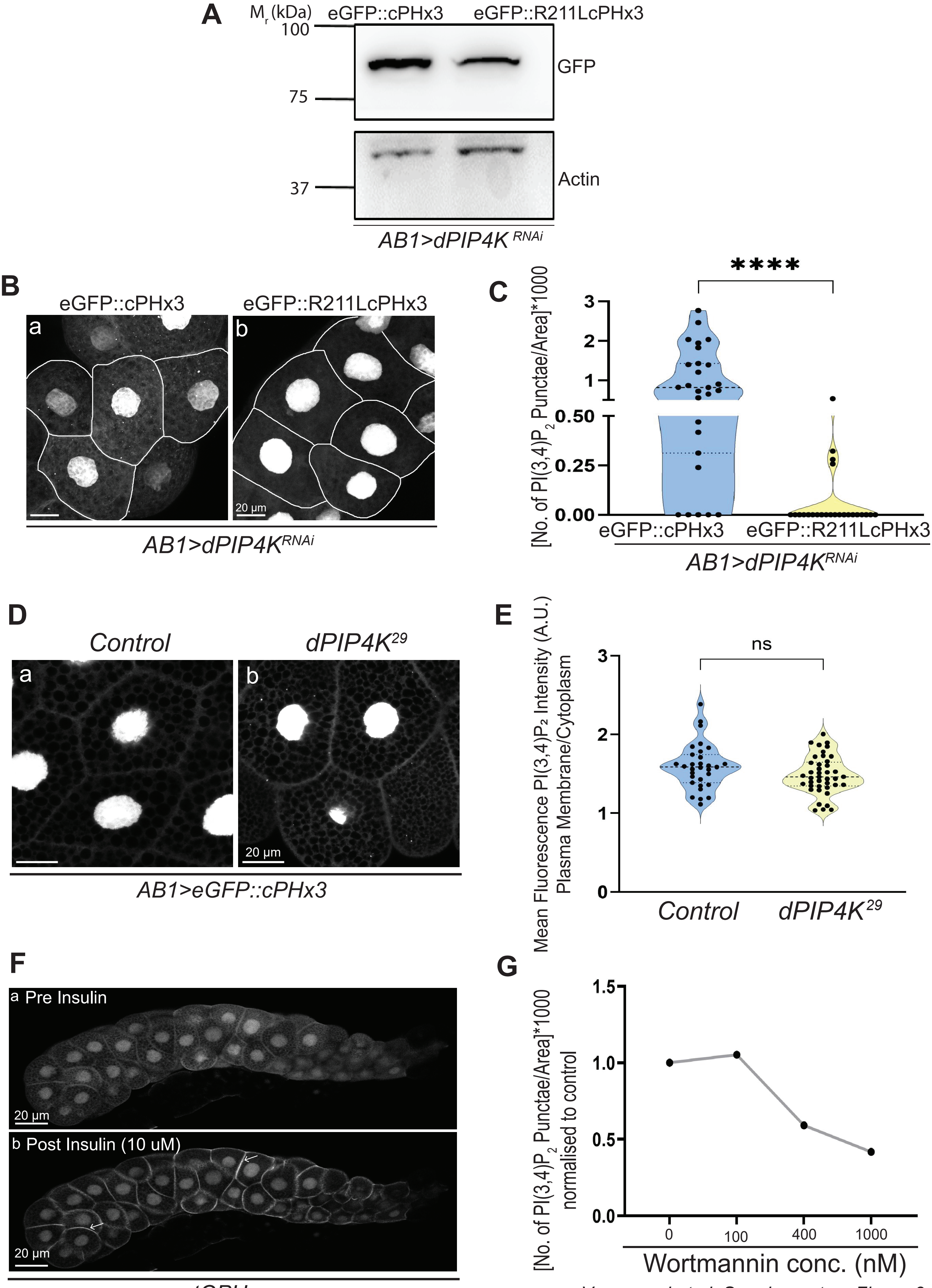


Venugopal et al.,Supplementary Figure 3

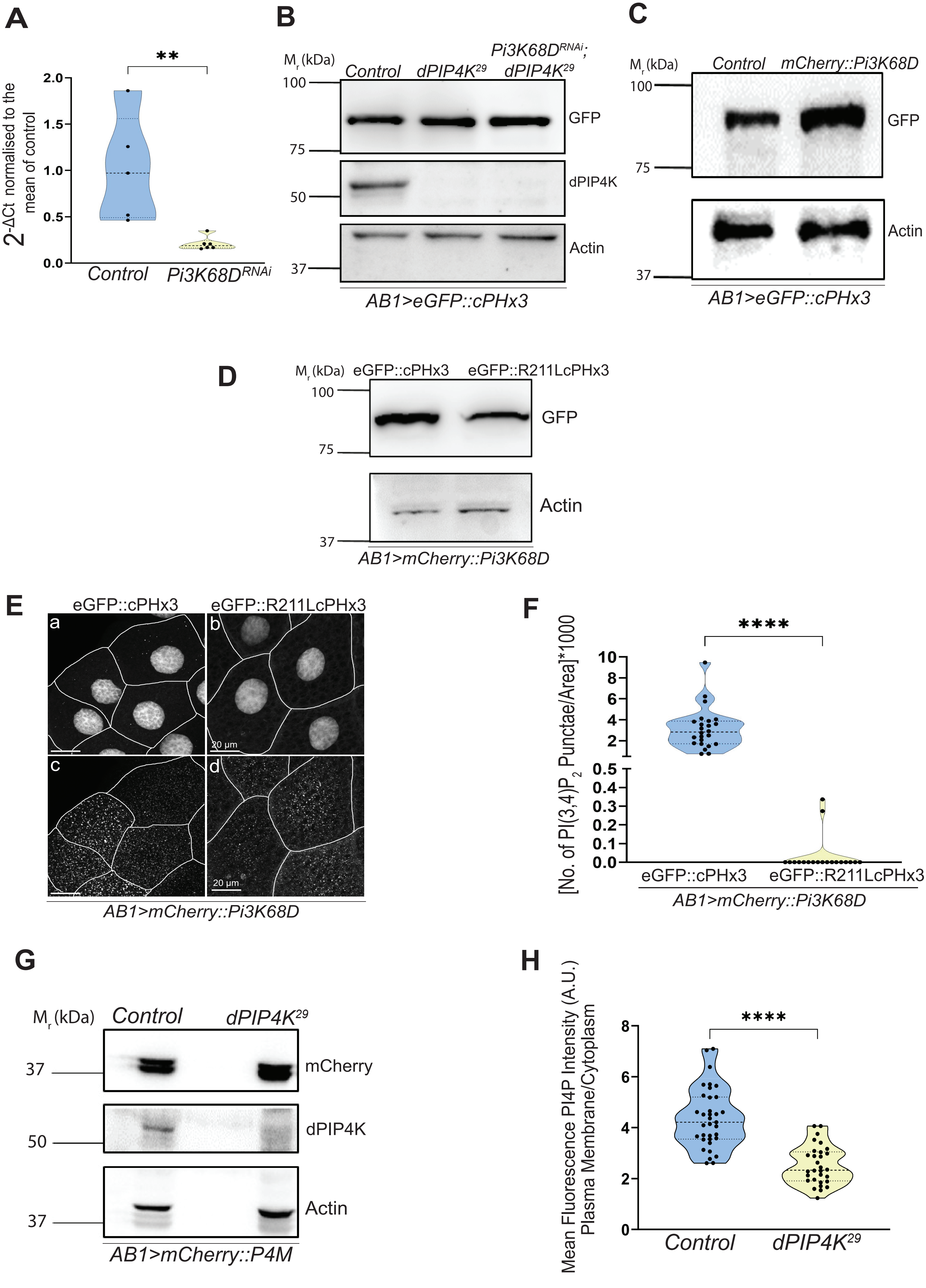


Venugopal et al.,Supplementary Figure 4

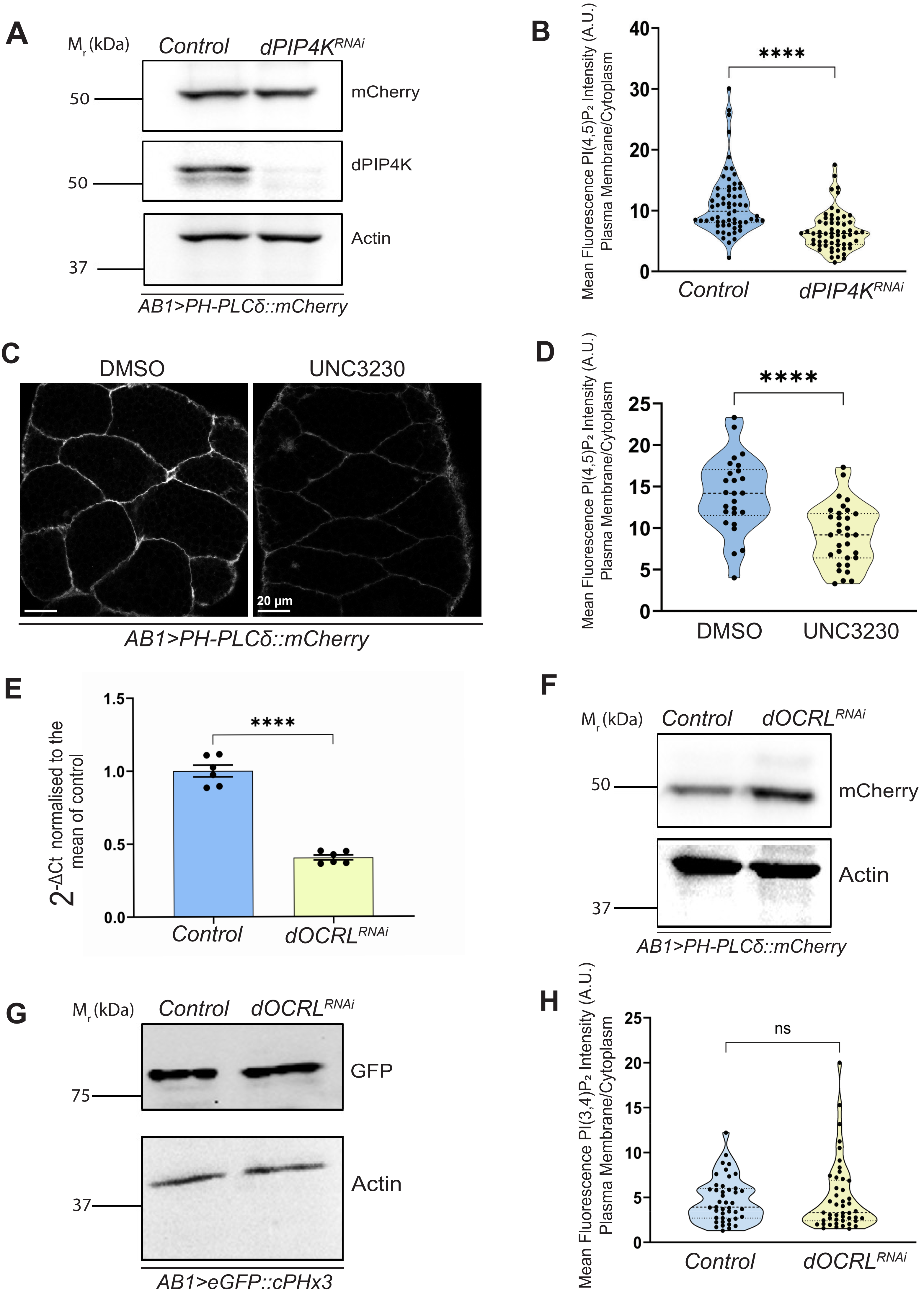


Venugopal et al.,Supplementary Figure 5
